# Trade-off shapes diversity in eco-evolutionary dynamics

**DOI:** 10.1101/184432

**Authors:** Farnoush Farahpour, Mohammadkarim Saeedghalati, Verena Brauer, Daniel Hoffmann

## Abstract

We introduce an Interaction and Trade-off based Eco-Evolutionary Model (ITEEM), in which species are competing for resources in a well-mixed system, and their evolution in interaction trait space is subject to a life-history trade-off between replication rate and competitive ability. We demonstrate that the strength of the trade-off has a fundamental impact on eco-evolutionary dynamics, as it imposes four phases of diversity, including a sharp phase transition. Despite its minimalism, ITEEM produces without further *ad hoc* features a remarkable range of observed patterns of eco-evolutionary dynamics. Most notably we find self-organization towards structured communities with high and sustainable diversity, in which competing species form interaction cycles similar to rock-paper-scissors games.

## I. INTRODUCTION

The mechanisms underlying emergence and stability of diversity in communities of organisms competing for resources have been debated for many years [1–5]. The competitive exclusion principle and the consumer-resource model theoretically restrict the number of coexisting species to the number of resources [6, 7] whereas we observe an immense diversity in natural communities [2, 4, 7] and controlled experiments [8–12] where many species share few resources. This apparent contradiction gives the stunning biodiversity in communities the air of a paradox [2, 13].

The situation becomes even more puzzling if we introduce evolution into ecological models as it should perturb ecological equilibria, select the fittest and discard the rest [14], and in this way also appears to reduce biodiversity. However on the other hand it is undeniable that evolution plays a fundamental role in phenotype adaptation and thus in the emergence of diversity [15–18].

To tackle this problem eco-evolutionary models have been equipped with features [5] like spatial structure [19–21], spatial and temporal heterogeneity [22–24], tailored interaction network topologies, e.g. by addition of cycles or modules [25–28], or by combining different types of interactions [29–32], and adjusted mutation-selection rate [33–35], to name but a few. However, it is unclear whether such features are necessary to explain biodiversity. For instance, we also observe diversity under stable and homogeneous conditions [9–12, 36]. Hence, there could still be a fundamental ingredient missing that could help to explain biodiversity more generally.

One candidate for such an ingredient are the life-history trade-offs. Their role for stabilizing diversity has been investigated in previous eco-evolutionary studies [37–41] and experiments [41–45]. It has been shown that if metabolic trade-offs are considered, the competitive exclusion principle need not to be observed in homogeneous environments [40, 46]. Trade-offs that regulate energy investment in different life-history strategies are imposed by physical laws such as energy conservation or other thermodynamic constraints [40, 42, 47]. These laws constrain evolutionary trajectories in trait space of species [48, 49], i.e. a species cannot adopt any combination of strategies at the same time.

In a system of evolving competitors, strategy adoption is the result of eco-evolutionary feedbacks mediated by interaction of species [31, 50–52]. Evolutionary changes at the genetic level influence ecology if they cause phenotypic variations that affect biotic or abiotic interactions of species which in turn changes the species composition and occasionally forces species to evolve their strategies [52–58]. This suggests that we do not need to follow all evolutionary changes at the genetic or phenotypic level, but only those that affect interactions [59–62]. Consequently, we have chosen a model in which interactions themselves are evolving. This neglect of genetic and phenotypic details equals a coarse-graining of the eco-evolutionary system, which not only reduces complexity but also should make the approach more widely applicable. It can also help us to study how complex competitive interaction networks evolve and shape diversity.

Here we introduce a new, minimalist model that incorporates the elements introduced above, the *Interaction and Trade-off based Eco-Evolutionary Model (ITEEM)*, to study the development of communities of organisms that compete for steadily supplied resources. It is a simple, intuitive eco-evolutionary model at the interaction level, and with a life-history trade-off between interaction traits and replication rate: better competitors replicate less [38, 63].

We show that ITEEM dynamics closely resembles observed eco-evolutionary dynamics, such as sympatric speciation [11, 64–66], emergence of two or more levels of differentiation similar to phylogenetic structures [67], large and complex biodiversity over long times [11, 12], evolutionary collapses and extinctions [27, 60], and emergence of cycles in interaction networks that facilitate species diversification and coexistence [28, 50, 68–71]. Interestingly, the model shows a unimodal (“humpback”) behavior of diversity as function of trade-off, with a critical trade-off at which biodiversity undergoes a phase transition, a behavior observed in nature [72–75]. ITEEM shows that diversity is a natural outcome of competition when species evolve under certain physical constraints which restrict the energy allocation in different strategies.

## II. MODEL

ITEEM is an individual-based model with simple intuitive updating rules for population and evolutionary dynamics. A simulated system in ITEEM has *N_s_* sites of undefined spatial arrangement (no neighborhood), each providing resources for metabolism of one organism with a constant rate. The community is well-mixed which means that the encountering probability for each pair of individuals is equal. We start an eco-evolutionary simulation with individuals of a single strain occupying a fraction of the *N_s_* sites and carry out long simulations for millions of generations. Note that in the following we use the term *strain* for a set of individuals with identical traits. In contrast, a *species* is a cluster of strains with some diversity (cluster algorithm described in Supplemental Material SM-1 [76]).

At every generation or time step *t*, we try *N_ind_(t)* (number of individuals at that time step) replications of randomly selected individuals. Each selected individual of a strain *α* can replicate with rate *r_α_*, with its offspring randomly mutated with rate μ to new strain *α′*. An individual will vanish if it has reached its lifespan, drawn at birth from a Poisson distribution with overall fixed mean lifespan λ. This is equivalent to an identical per capita death rate for all species.

Each newborn individual is assigned to a randomly selected site. If the site is empty, the new individual will occupy it. If the site is already occupied, the new individual competes with the current holder in a life-or-death struggle; In that case, the surviving individual is determined probabilistically by the “interaction” *I_αβ_*, defined for each pair of strains α, β. *I_αβ_* is the survival probability of an α individual in a competitive encounter with a β individual, with *I_αβ_* ∈ [0, 1] and *I_αβ_* + *I_βα_* = 1. All interactions *I_αβ_* form an interaction matrix **I**(*t*) that encodes the outcomes of all possible competitive encounters.

If strain α goes extinct, the αth row and column of **I** are deleted. Conversely, if a mutation of α generates a new strain α′, **I** grows by one row and column:

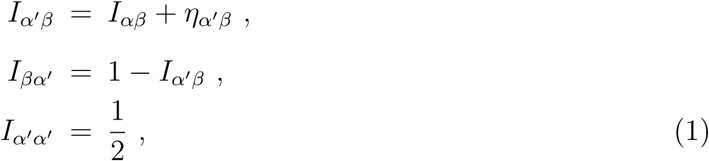

where α′ inherits interactions from α, but with small random modification ηα′β, drawn from a zero-centered normal distribution of fixed width *m*. Rowα of **I** can be considered the “interaction trait” **T**_α_ = (*I_α1_, I_α2_, …, I_α*N_sp_(t)*_) of strain α, with *N_sp_(t)* the number of strains at time *t*. Evolutionary variation of mutants in ITEEM can represent any phenotypic variation which influences direct interaction of species and their relative competitive abilities [77–80]*.

To implement trade-offs between fecundity and competitive ability, we introduce a relation between replication rate *r_α_* (for fecundity) and competitive ability *C*, defined as average interaction

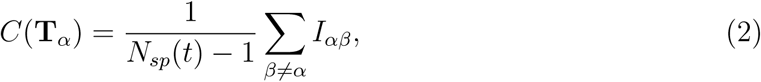

and we let this relation vary with trade-off parameter *s*:

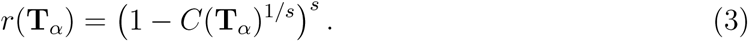

With Eq. 3 better competitive ability leads to lower fecundity and vice versa. Of course, other functional forms are conceivable. To systematically study effects of trade-off on dynamics we varied 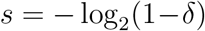 with *trade-off strength ö* covering [0,1] in equidistant steps (SM-2). The larger *δ*, the stronger the trade-off. *δ = 0* makes *r =1* and thus independent of *C* which means no trade-off.

We compare ITEEM results to the corresponding results of a neutral model [81], where we have formally evolving vectors **T**_α_, but fixed and uniform replication rates and interactions. Accordingly, the neutral model has no trade-off.

ITEEM belongs to the well-established class of generalized Lotka-Volterra models in the sense that the mean-field version of our stochastic, individual-based model leads to competitive Lotka-Volterra equations (SM-3).

## III. RESULTS

### Generation of diversity

Our first question was whether ITEEM is able to generate and sustain diversity. Since we have a well-mixed system with initially only one strain, a positive answer implies sympatric diversification, i.e. emergence of new strains and species via evolution without geographic isolation or resource partitioning. In fact, we observe that during long-time eco-evolutionary trajectories in ITEEM, new distinct species emerge and their coexistence establishes a sustainable high diversity in the system (Fig. 1a). Remarkably, the emerging diversity has a clear hierarchical cluster structure (Fig. 1b): at the highest level we see well-separated clusters in trait space similar to biological *species*. Within these clusters there are sub-clusters of individual strains (SM-4) [67]. Both levels of diversity can be quantitatively identified as levels in the distribution of branch lengths in minimum spanning trees in trait space (SM-5). This hierarchical diversity is reminiscent of the phylogenetic structures in biology [67]. Overall, the model shows evolutionary divergence from one ancestor to several species consisting of a total of hundreds of co-existing strains (Fig. 1c and supplementary animation). In the course of this diverging sympatric evolution diversity measures like richness, entropy, or functional diversity typically increase and, depending on trade-off parameter δ, this high diversity can be sustained over hundreds of thousands of generations (Fig. 1d, and SM-6). Collapses to low diversity occur rarely and are usually followed by recovery of diversity.

**Figure 1.**
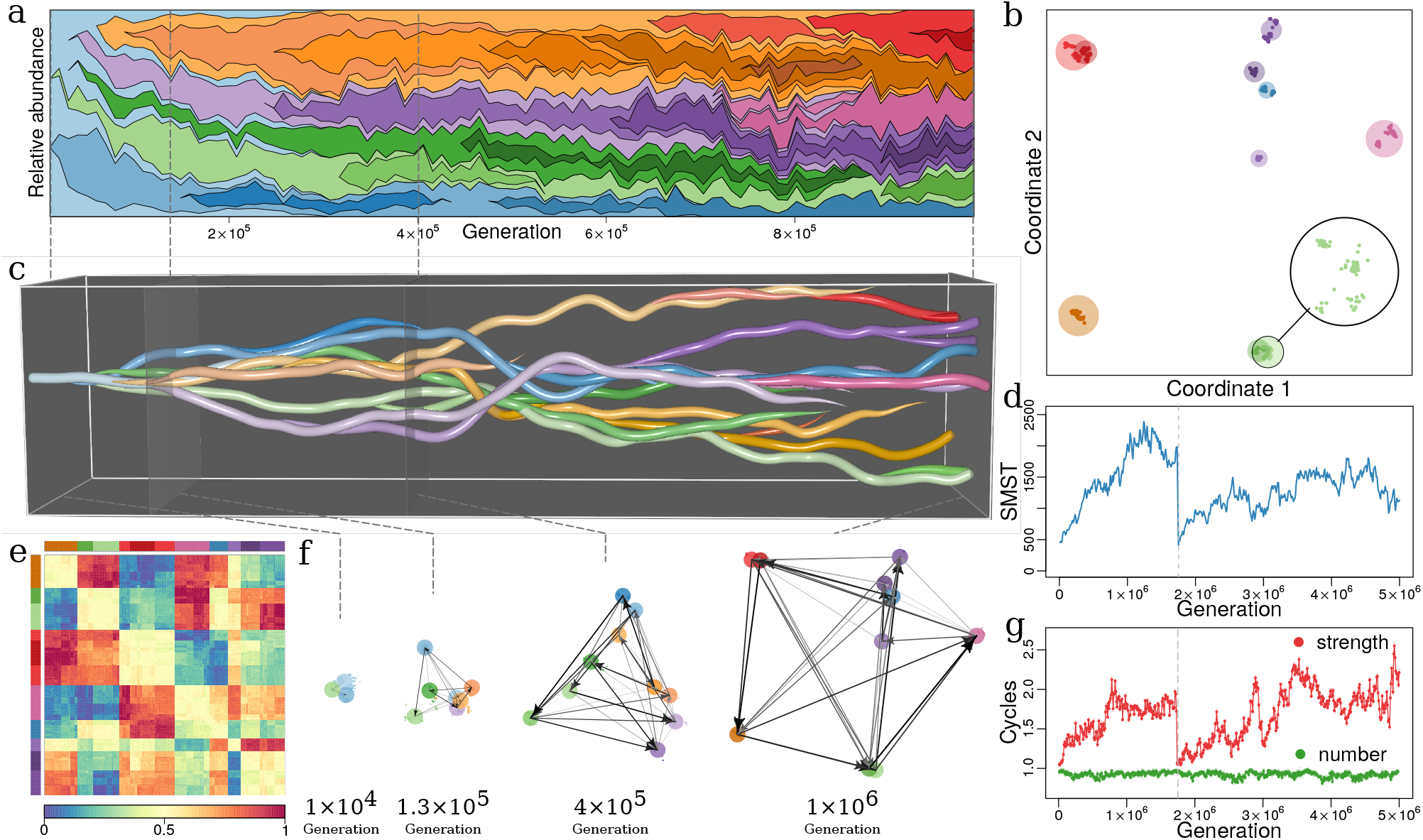
Evolutionary dynamics of a community driven by competitive interactions, with trade-off between fecundity and competitive abilities (*δ = 0.5, λ = 300, μ = 0.001, m = 0.02, N_s_* = 10^5^). (a) **Species’ frequencies over time** (Muller plot): one color per species, vertical width of each colored region is relative abundance of respective species. Frequencies are recorded every 10^4^ generations over 10^6^ generations. The plot is produced using MullerPlot package in R. (b) **Distribution over trait space**: Snapshot of distribution in trait space after 10^6^ generations, reduced to two dimensions that explain most of the variance in trait space (see SM-4). Points and discs are strains and species, respectively (see SM-1). Magnified disc in lower right corner shows strains in the light green species disc. Disc diameter scales with abundance of species. This snapshot consists of 660 strains in 10 species. (c) **Evolutionary dynamics in trait space**: Snapshots as in panel (b), but concatenated for all times (horizontal axis), from the monomorphic first generation to 10^6^ generation. (d) **Functional diversity over time** in terms of the size of minimum spanning tree (SMST) in trait space (SM-6). At 1.75 × 10^6^ generations an evolutionary collapse happens in which all species but one go extinct (vertical dashed line). (e) **Heatmap of interaction matrix I** for generation 10^6^. Row and column order reflects species clusters, consistent with panel (b) and indicated by color bars along top and left. Colors inside heatmap represent interaction rates (color-key along bottom). (f) **Evolution of interaction network**: several snapshots from panel (c) with interactions between species (colored discs) as directed edges. Directions and strengths of edges given by signs and absolute values, respectively, of averages over ***I**_αβ_ − **I**_βα_*, with *α, β* the component strains of the species linked by edge. (g) **Numbers and average strengths of cycles over time** in green and red, respectively. The strength of a cycle is defined by its weakest edge. Number and average strength given in units of number and average strength of equivalent random network, respectively (SM-7). Right ends in (a) and (c) correspond to panel (b) and (e). Colors of species are the same in panels (a), (b), (c), (e) and (f). Note that time scales differ between panels (a), (c) and (d), (g).

The observed divergence contradicts the long-held view of sequential fixation in asexual populations [82]. Instead, we see frequently concurrent speciation with emergence of two or more species in quick succession (Fig. 1a), in agreement with recent results from long-term bacterial and yeast cultures [11, 12, 83].

ITEEM systems self-organize towards structured communities: the interaction matrix of a diverse system obtained after many generations shows groups of strains with similar interaction strategies, and these groups are well-separated from other such groups (Fig. 1b, e) [84]. Interaction trait in ITEEM directly determines the functionality of strains and species in their community and these groups. Fig. 1e shows self-organization of well-separated functional niches [85–87] each occupied by a group of closely related strains. This niche differentiation among species which facilitates the possibility of coexistence is the result of frequency-dependent selection among competing strategies. Within each functional niche, occupied by strains with similar strategies, the predominant dynamics determining their relative abundances is neutral. Subsequent speciation can occur in each group when genetic drift makes a sufficiently large difference in their strategies that boosts selection forces.

Our model also allows to study speciation in detail, e.g. in terms of interaction network dynamics. The interaction matrix **I** defines a complete graph, and we determined direction and strength of interaction edges between two strains *α, β* as sign and size of *I_αβ_ − I_βα_*, respectively. Accordingly, for the interaction network of *species* (i.e. clusters of strains) we computed directed edges between any two species by averaging over inter-cluster edges between the strains in these clusters. The strength and direction of interaction edges marks the effective flow of population between species. We observe in ITEEM simulations that strong edges appear in the interaction network as the trait distance (or dissimilarity) of species increases (Fig. 1f) [88].

Three or more directed edges can form cycles of strains in which each strain competes successfully against one cycle neighbor but loses against the other neighbor, a configuration corresponding to rock-paper-scissors games [89]. Such intransitive interactions have been observed in nature [79, 90–92], and it has been shown that they stabilize a system driven by competitive interactions [20, 50, 68, 70]. Here, we find that the increase of diversity coincides with growth of average cycle strength (Figs 1d, g and SM-7). Note that these cycles emerge and self-organize in the evolving ITEEM networks without any presumption or constraint on network topology.

### Impact of trade-off and lifespan on diversity

The eco-evolutionary dynamics described above depend on lifespan and trade-off between replication and competitive ability. To show this we study properties of interaction matrix and trait diversity. Fig. 2a relates average interaction rate 〈*I*〉 and average cycle strength ρ to trade-off parameter δ at fixed lifespan λ. Fig. 2b summarizes the behavior of diversity as function of δ and λ. Overall, we see in this phase diagram a weak dependency on λ and a strong impact of δ, with four distinct phases (I-IV) from low to high δ.

**Figure 2.**
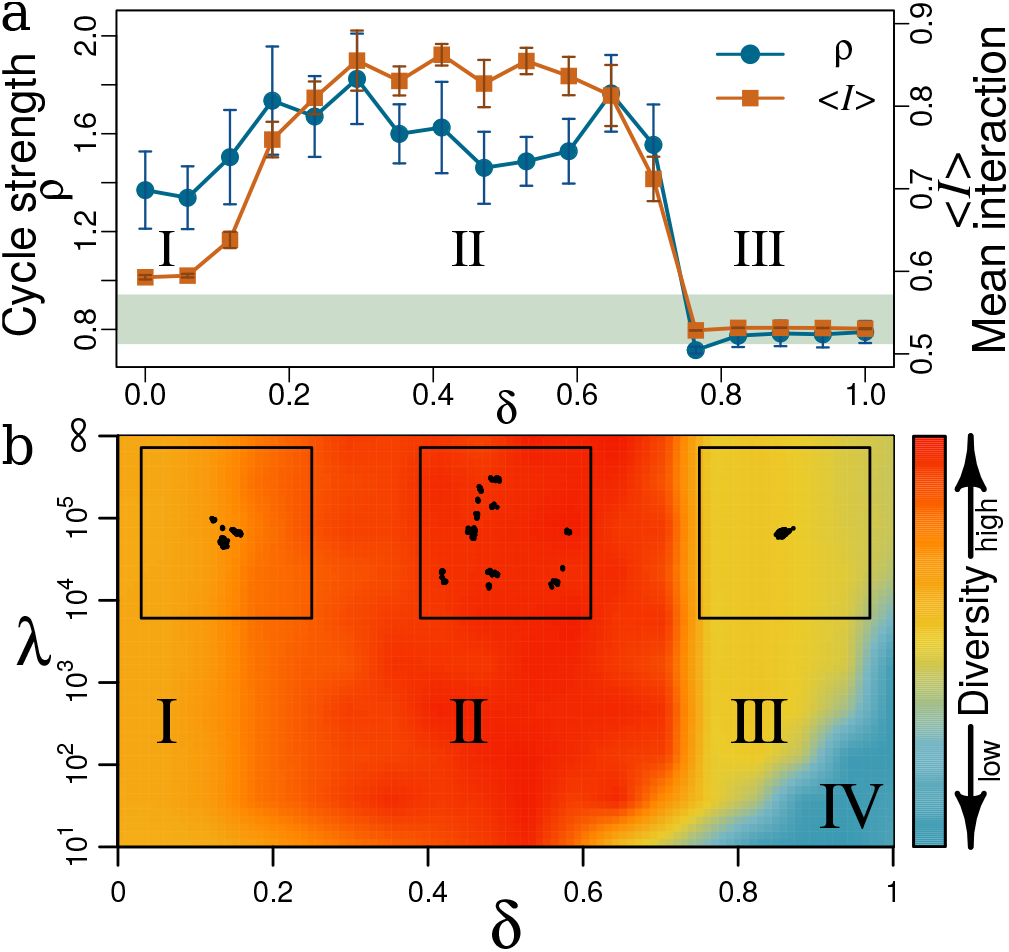
Effects of trade-off δ and lifespan λ on community structure and diversity. (a) Average interaction rates *〈I〉* (orange squares) and average strength of cycles ρ (blue circles) as function of δ. Average cycle strength is given in units of average strength of random networks for the respective trade-off (SM-7). Averages are calculated over three different simulations, each over 5 × 10^6^generations with *μ = 0.001, m = 0.02, λ = ∞ and N_s_ = 10^5^*. Error bars are standard deviations averaged over three concatenated simulations. The shaded area marks cycle strength for a neutral model with corresponding parameters ± standard deviation. (b) Phase diagram of diversity as function of *δ* and *λ*. Diversity is given as consensus of several quantities (SM-6). Four phases (I-IV) can be distinguished. Insets at the top margin are representative MDS plots (SM-4) of strain distributions in trait space, as in Fig. 1b, with λ = 10^5^ but different values of Ô (left to right: I with δ = 0.11; II with δ = 0.5; III with δ = 0.89). Panel (a) corresponds to a horizontal cross-section through the phase diagram in panel (b) with λ = ∞ for 〈*I*〉 and ρ as diversity measures.

Without trade-off, strains do not have to sacrifice replication rate for better competitive abilities. We have a low-diversity population dominated by Darwinian demons, species with high competitive ability and replication rate. Quick predominance of such strategies impedes formation of a diverse network. Increasing δ in phase I (0 <δ ≲ 0.2) slightly increases 〈*I*〉 and ρ (Fig. 2a): biotic selection pressure exerted by inter-species interactions starts to generate diverse communities (left inset in Fig. 2b, SM-6). However, the weak trade-off still favors investing in higher competitive ability. When increasing δ further (phase II), tradeoff starts to force strains to choose between higher replication rate *r* or better competitive abilities *C*. Neither extreme generates viable species: sacrificing *r* completely for maximum *C* stalls species dynamics, whereas maximum *r* leads to inferior *C*. Thus strains seek middle ground values in both *r* and *C*. The nature of *C* as mean of interactions (Eq. 2) allows for many combinations of interaction traits with approximately the same mean. Thus in a middle range of *r* and *C*, many strategies with the same overall fitness are possible, which is a condition of diversity [93]. From this multitude of strategies, sets of trait combinations emerge in which strains with different combinations keep each other in check, e.g. in the form of competitive rock-paper-scissors-like cycles between species described above. An equivalent interpretation is the emergence of diverse sets of non-overlapping compartments or trait space niches (Fig. 1b,e). Diversity in this phase II is the highest and most stable (middle inset in Fig. 2b, SM-6). As δ approaches 0.7, *〈I〉* and ρ plummet (Fig. 2a) to interaction rates comparable to noise level *m*, and a cycle strength typical for the neutral model (horizontal gray ribbon in Fig. 2a), respectively. The sharp drop of 〈*I*〉 and ρ at δ ≈ 0.7 is reminiscent of a phase transition. As expected for a phase transition, the steepness increases with system size (SM-8). For δ ≳ 0.7 interaction rates never grow and no structure emerges; diversity remains low and close to a neutral system. The sharp transition at δ ≈ 0.7 which is visible in practically all diversity measures (between phases II and III in Fig. 2b, SM-6) is a transition from a system dominated by biotic selection pressure to a neutral system. In high-trade-off phase III, any small change in *C* changes *r* drastically. For instance, given a strain *S* with *r* and *C*, a closely related mutant *S’* with 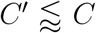 will have *r’≫ r* (because of the large trade-off), and therefore will invade *S* quickly. Thus, diversity in phase III will remain stable and low, characterized by a group of similar strains with no effective interaction and hence no diversification to distinct species (right inset in Fig. 2b, SM-6). In this high trade-off regime, lifespan comes into play: here, decreasing λ can make lives too short for replication. These hostile conditions minimize diversity and favor extinction (phase IV).

### Trade-off, resource availability, and diversity

There is a well-known but not well understood unimodal relationship between biomass productivity and diversity (“humpback curve”,[73, 74]): diversity culminates once at middle values of productivity. This behavior is reminiscent of horizontal sections through the phase diagram in Fig. 2b, though here the driving parameter is not productivity but trade-off. However, we can make the following argument for a monotonous relation between productivity and trade-off. First we note that biomass productivity is a function of available resources: the larger the available resources, the higher the productivity. This allows us to argue in terms of available resources. If then a species has a high replication rate in an environment with scarce resources, its individuals will not be very competitive since for each of the numerous offspring individuals there is little material or energy available. On the other hand, if a species under these resource-limited conditions has competitively constructed individuals it cannot produce many of them. This corresponds to a strong trade-off between replication and competitive ability for scarce resources. At the opposite, rich end of the resource scale, species are not confronted with such hard choices between replication rate and competitive ability, i.e. we have a weak trade-off. Taken together, the trade-off axis should roughly correspond to the inverted resource axis: strong trade-off for poor resources (or low productivity), weak trade-off for rich resources (or high productivity); a detailed analytical derivation will be presented elsewhere. The fact that ITEEM produces this humpback curve that is frequently observed in planktonic systems[74] proposes trade-off as underlying mechanism of this productivity-diversity relation.

### Frequency-dependent selection

Observation of eco-evolutionary trajectories as in Fig. 1 suggested the hypothesis that speciation events in ITEEM simulations do not occur with a constant rate and independently of each other, but that one speciation makes a following speciation more likely. We therefore tested whether the distribution of time between speciation or extinction events is compatible with a constant rate Poisson process (SM-9). At long inter-event times we see the same decaying distribution for the Poisson process and for the ITEEM data. However, for shorter times there are significant deviations from a Poisson process for speciation and extinction events: at inter-event times of around 10^4^ the number of events *decreases* for a Poisson process but significantly *increases* in ITEEM simulations. This confirms the above hypothesis that emergence of new species increase the probability for generation of further species, and additionally that loss of a species makes further losses more likely. This result is similar to the frequency-dependent selection observed in microbial systems where new species open new niches for further species, or the loss of species causes the loss of dependent species [11, 83].

The above analysis suggests a further application of ITEEM simulations. Eco-evolutionary trajectories from ITEEM simulations can be used to develop analytical methods for the inference of competition based on observed diversification patterns. Such methods could be instrumental for understanding the reciprocal effect of competition and diversification.

### Effect of mutation rate on diversity

Simulations with different mutation rates (μ = 10^−4^, 5 × 10^−4^,10^−3^, 5 × 10^−3^) show that in ITEEM diversity grows faster and to a higher level with increasing mutation rate, but without changing the overall structure of the phase diagram (SM-10). One interesting tendency is that for higher mutation rates, the lifespan becomes more important at the interface of regions III and IV (high trade-offs), leading to an expansion of region III at the expense of hostile region IV: long lifespans in combination with high mutation rate establish low but viable diversity at strong trade-offs. The humpback curve of diversity over trade-off strength is observed for all mutation rates. Thus, the diversity in ITEEM is not a simple result of a mutation-selection balance but it emphasizes that trade-off plays an important role in expanding diversity in trait space.

### Comparison of ITEEM with neutral model

The neutral model introduced in the Model section has no meaningful interaction traits, and consequently no meaningful competitive ability or trade-off with replication rate. Instead, it evolves solely by random drift in phenotype space. Similarly to ITEEM, the neutral model generates a clumpy structure in trait space (SM-11), though here the clusters are much closer and thus the functional diversity much lower. This can be demonstrated quantitatively by the size of the minimum spanning tree of populations in trait space that are much smaller for the neutral model than for ITEEM at moderate trade-off (SM-11). The clumpy structures generated in neutral model do not follow a stable trajectory of divergent evolution and hence niche differentiation can not be established. In a neutral model, without presence of frequency-dependent selection and trade-off, stable structures and cycles can not be constructed in the interaction network, and consequently diversity does not grow effectively (SM-11). The comparison with the neutral model points to frequency-dependent selection as a promoter of diversity in ITEEM. For high trade-offs (region III, Fig. 2b), diversity and number of strong cycles in ITEEM are comparable to the neutral model (Fig. 2a.)

## IV. DISCUSSION

Interaction based eco-evolutionary models have received some attention in the past [59–62] but then were almost forgotten, despite remarkable results. We think that these works have pointed to a possible solution of a hard problem: The complexity of evolving ecosystems is immense, and it is therefore difficult to find a representation suitable for the development of a statistical mechanics that enables qualitative and quantitative analysis [52]. Modeling in terms of interaction traits, rather than detailed descriptions of genotypes or phenotypes, then coarse-grains these complex systems in a natural, biologically meaningful way.

Despite these advantages, interaction based models so far have not shown some key features of real systems, e.g. emergence of large, stable and complex diversity, or mass extinctions with the subsequent recovery of diversity [27, 61]. Therefore, interaction based models were supplemented by *ad hoc* features, such as special types of mutations [61], induced extinctions [94], or enforcement of partially connected interaction graphs [27].

An essential component missing in previous models with evolving interactions had been a constraint on strategy adoption. In real systems such constraints prevent the emergence of Darwinian demons, i.e. species that in the absence of any restriction develop and act as a sink in the network of population flow. Life-history trade-offs, like the trade-off between replication and competitive ability that have now been experimentally established as essential to living systems [42, 44], are inescapable constraints imposed by physical limitations in natural systems. Our results with ITEEM show that trade-offs fundamentally impact eco-evolutionary dynamics, in agreement with other eco-evolutionary models with trade-off [38, 39, 46, 95]. Remarkably, we observe with ITEEM sustained high diversity in a well-mixed homogeneous system, without violating the competitive exclusion principle. This is possible because moderate life-history trade-offs force evolving species to adopt different strategies or, in other words, lead to the emergence of well-separated functional niches in interaction space [40, 46].

Given the accumulating experimental and theoretical evidences, the importance of tradeoff for diversity is becoming more and more clear. ITEEM provides an intuitive and generic conceptual framework with a minimum of specific assumptions or requirements. This makes the results transferable to different systems, e.g. biological, economical and social systems, wherever competition is the driving force of evolving community. Put simply, ITEEM shows generally that in a simple eco-evolutionary model with of a standard population dynamics (birth-death-competition) and a basic evolutionary process (mutation), diverse set of strategies will emerge and coexist if species are forced to manage their resource allocation.

Despite its simplifications, ITEEM reproduces in a single framework several phenomena of eco-evolutionary dynamics that previously were addressed with a range of distinct models or not at all, namely sympatric and concurrent speciation with the emergence of new niches in the community, recovery after mass-extinctions, large and sustained functional diversity with hierarchical organization, spontaneous emergence of intransitive interactions and cycles, and a unimodal diversity distribution as function of trade-off between replication and competition. The model allows detailed analysis of mechanisms and could guide experimental tests.

The current model has important limitations. For instance, the trade-off formulation was chosen to reflect reasonable properties in a minimalistic way, that should be revised or refined as more experimental data become available. Secondly, individual life-spans come from a random distribution with an identical fixed mean. Hence we have no adaptation and evolutionary-based diversity in life-span. This limits the applicability of the model just to the communities of species which are closely related and those which put their major adaptation effort in growth or reproduction rate and competitive ability. Furthermore, we have assumed a single, limiting resource in a well-mixed system to investigate the mechanisms behind diversification in competitive communities and possibility of niche differentiation without resource partitioning or geographic isolation. However, in nature, there will in general be several limiting resources and abiotic factors. It is possible to consider those as additional rows and columns in the interaction matrix I and including abiotic interactions as well as biotic ones.

## Acknowledgments

We thank S. Moghimi-Araghi for helpful suggestions on trade-off function.

## Supplementary Information

### I Species and strains

In literature there is no unique definition of species [1]. The ambiguity of the species concept is more severe in asexual population [1,2]. Here we use phylogenetic tree to define and classify species based on the point of divergence from a common ancestor (See [2] for discussion on pros and cons of this definition). Each branch rooted from a point of divergence in phylogenetic tree is counted as an individual species when it has endured more than 5 × 10^4^ generations considering all of its sub branches. Otherwise they are counted as the strains of their parents.

### II Trade-off function

Trade-off function used in this paper (Eq. 2 in manuscript) maps competitive ability of species to a replication rate in the range [0,1]. Its functionality (by changing the exponent) allows us to study effect of the form of trade-off (convex,concave and linear) on evolutionary process. As it is described in the manuscript, to systematically study effects of trade-off on dynamics we varied 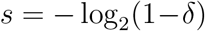 with *trade-off strength δ* covering [0,1] in equidistant steps. The larger δ, the stronger the trade-off. δ = 0 makes *r = 1* and thus independent of *C* which means no trade-off.

### III Generalized Lotka-Volterra (GLV) equation

The mean-field equation equivalent to the population dynamics of the agent-based model introduced in the manuscript is a competitive generalized Lotka-Volterra (GLV) equation. To show this we start from the main equation of population dynamics for our model:

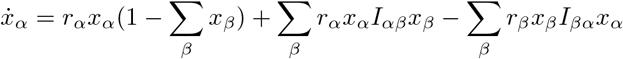

In which 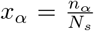 is the relative abundance or probability of finding species **α** in the system. The first term in this equation shows the increase in population of species **α** when it produces an offspring and is able to find an empty space in the system. The second term shows the increase in population of **α** when after reproduction, its offspring is able to invade another species and the last term shows the decrease in its population due to invasion of offspring of other species. We can rewrite the equation as follows:

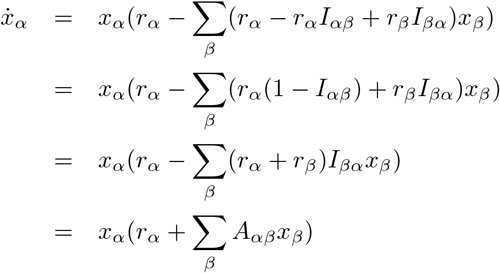

In which we used 1 − *I_αβ_ = I_βα_*(see Eq. 1 in manuscript). The last equation, 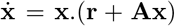, is the GLV equation. **A** is the community matrix and its elements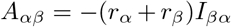 are always negative which means that our species are in direct competition with each other. In this derivation we didn’t consider the death rate or attributed lifespan of species and it can fit best with the result of **λ** = ∞

Equivalence of the agent-based model with GLV equation shows clearly that species are competing in a sense that they change the population of other competitors in order to obtain the resource in favor of their species fitness.

### IV Classical multi-dimensional scaling (CMDS)

The goal of an MDS algorithm is to put each object of a group in N-dimensional space such that the between-object distances are preserved as well as possible. They are usually used in information visualization, in particular to display the information contained in a distance matrix. The decision about number of dimensions can be done by looking at the eigenvalues of factor analysis.

We calculate the distance between all trait vectors **T**_α_ = (*I_α_,I_α2_,…, I_αN_sp_(t)_*) and by applying *cmdscale* function of R package (version 3.3.0) to the obtained distance matrix we project our trait space into 2D plots. The distance measure is Euclidean. Each point in the cMDS plot represents a trait vector, i.e. a strain. The larger the distance between two points, the more different the trait of corresponding species. cMDS is done merely to make it visually easier to see the evolutionary process in trait space. All other analysis are done directly in the original high-dimensional trait space.

In very early stage of evolution (Fig. 2-top) that species are very similar, 2 dimension is not sufficient to project them but when speciation and evolutionary speciation and branching occur (Fig. 2-bottom) it is more and more likely to project the trait space to 2D. This is clear by looking at eigenvalues of factor analysis. Fig. 3 shows Percentage of Variation Explained by the first two coordinates versus time for one simulation.

### V Distribution of species and strains in trait space

One way to see how species and strains are distributed over trait space is to look at the minimum spanning tree (MST) of traits in trait space. For a hierarchical structure, big edges appear between clusters or modules and small edges connect the nodes inside each cluster (Fig. 4-left). We plotted the lengths of the sorted edges of MST versus their ranks (Fig. 4-right) for 500 snapshot of simulations with *δ = 0.5, λ = 300, μ = 0.001 and m = 0.02* From this plot we see that there is two clearly different scales in the size (Note the logarithmic scale of the plot). *R_1_* is a representative value for the size of clusters (distance between strains of one typical species) and R_2_ is a representative value for the size of the trait space (typical distance between different species). *N* could be considered as a representative value for the number of distinct clusters (species) in the system.

### VI Diversity indexes and parameters of dynamics

No Diversity index is sufficient to describe complete characteristics of a community. For example richness, i. e. number of species, has no information about the distribution of population among species. Evenness or equivalently Shannon entropy takes into account this distribution but has no information about the diversity of the trait of species, i.e. how diverse is a system in functionality of its species. Functional diversity indexes focus on this aspect but none of them is comprehensive to comprise all information about properties of trait space [3]. Information about density of species over resources, rate of extinction and emergence and interaction of species should be also considered to assay the dynamics of a community comprehensively.

#### A Diversity over time

Next plots show how a typical diversity index changes over time. Note that evolutionary collapses (mass extinctions) occasionally occur.

Fig. 5 shows one of the indexes (Size of Minimum Spanning Tree (SMST)) over time for different tradeoffs but same lifespan in each plot.Fig. 6 shows the same index (SMST) over time for different lifespans but the same trade-off in each plot.

#### B Diversity indexes and parameters of dynamics for different trade-offs and lifespans

In Fig. 2 of the manuscript we have used normalized values of “richness”, “Shannon entropy”, “standard deviation of reproduction rate”, “maximum distance in trait space”, “standard deviation of interaction rates”, “sum of squared lengths of minimum spanning tree of trait space”, “functional diversity indexes, i.e functional dispersion, Rao index and functional evenness (all in two versions: with and without abundance)” and “strength of cycles” to compute an average as a descriptive dimensionless parameter for different regions. Although that plot can give a general idea about different behavior of communities with different δ and λ, but details of diversity indexes must be considered carefully. In Fig. 7 some of the most important parameters measured in our simulations to evaluate the community state are reported. Plots show the parameters for different trade-offs and lifespans. The Index values are averaged over 5 × 10^6^ Generations.

### VII Cycles

Local flow of energy/mass in our community is determined by matrix ***I**_αβ_ − **I**_βα_* which is an asymmetric matrix and its corresponding network is a directed network with one directed edge between each pair of nodes with weight between 0 and 1. We compare number and strength of cycles of this evolving network with its equivalent shuffled random networks. For this purpose we need

1. a method to find a cycle: We select 3 nodes of the network randomly and check if they form a cycle of size 3. If yes, it increases the number of cycles by 1 and the minimum weight of its edges (limiting path in energy/mass flow) would be assigned as its strength. This procedure is repeated many times (> 10^5^) and then we average over the strengths.
2. to build random networks: We shuffle the edges of original network to obtain a random network. Then we apply the procedure described in 1 on this network to measure the number of cycles and their strengths. This step is done several times (> 10) to have different random networks.

Result of these two steps are plotted in Fig. 1-g and Fig. 2-a in the paper.

**Figure 1.**
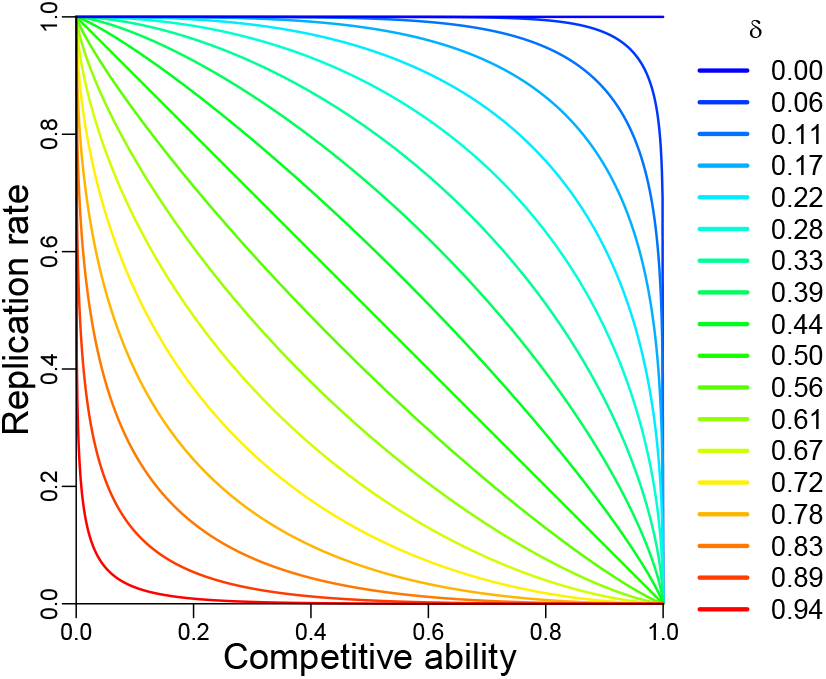
Trade-off functions

**Figure 2.**
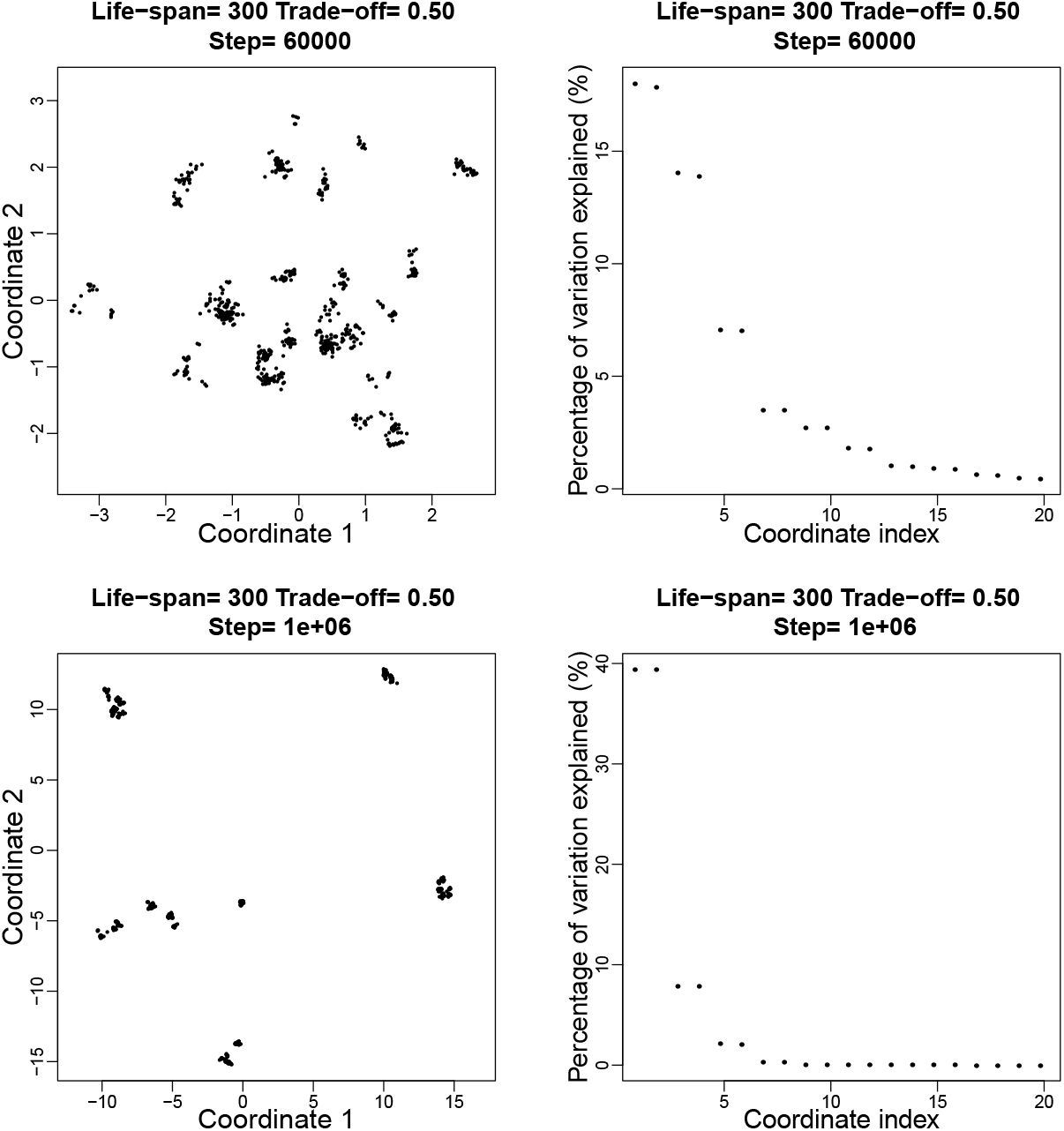
Top-left: Trait space at generation 6 × 10^4^ projected into 2 dimensions using CMDS. Top-right: Percentage of variation explained by eigenvectors (correspond to the first 20 eigenvalues) of factor analysis. Here the first two eigenvectors explain around 36% of variation. Bottom: The same as the top but for generation 1 × 10^6^. Here the first two eigenvectors explain around 78% of variation. Simulation is done with *δ = 0.5, λ = 300, μ = 0.001 and m = 0.02*

**Figure 3.**
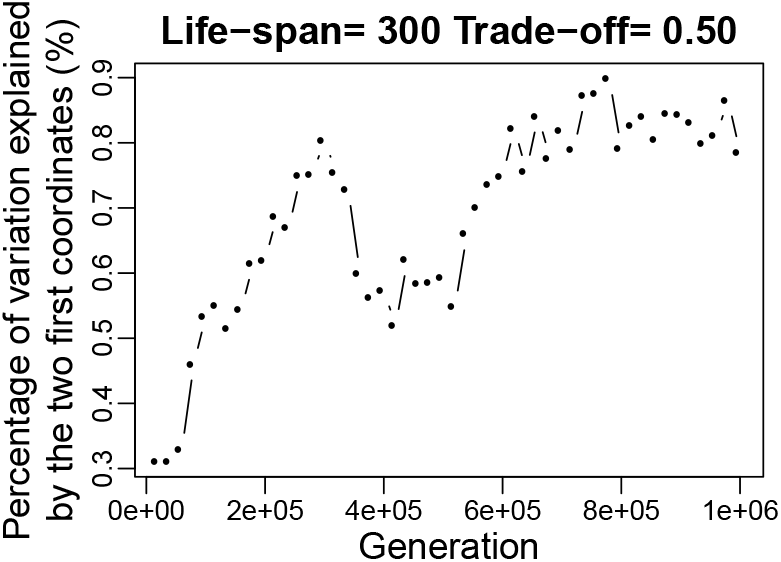
Percentage of variation explained by the first two coordinates versus time for one simulation. Simulation is done with *δ = 0.5, λ = 300, μ = 0.001 and m = 0.02*

**Figure 4.**
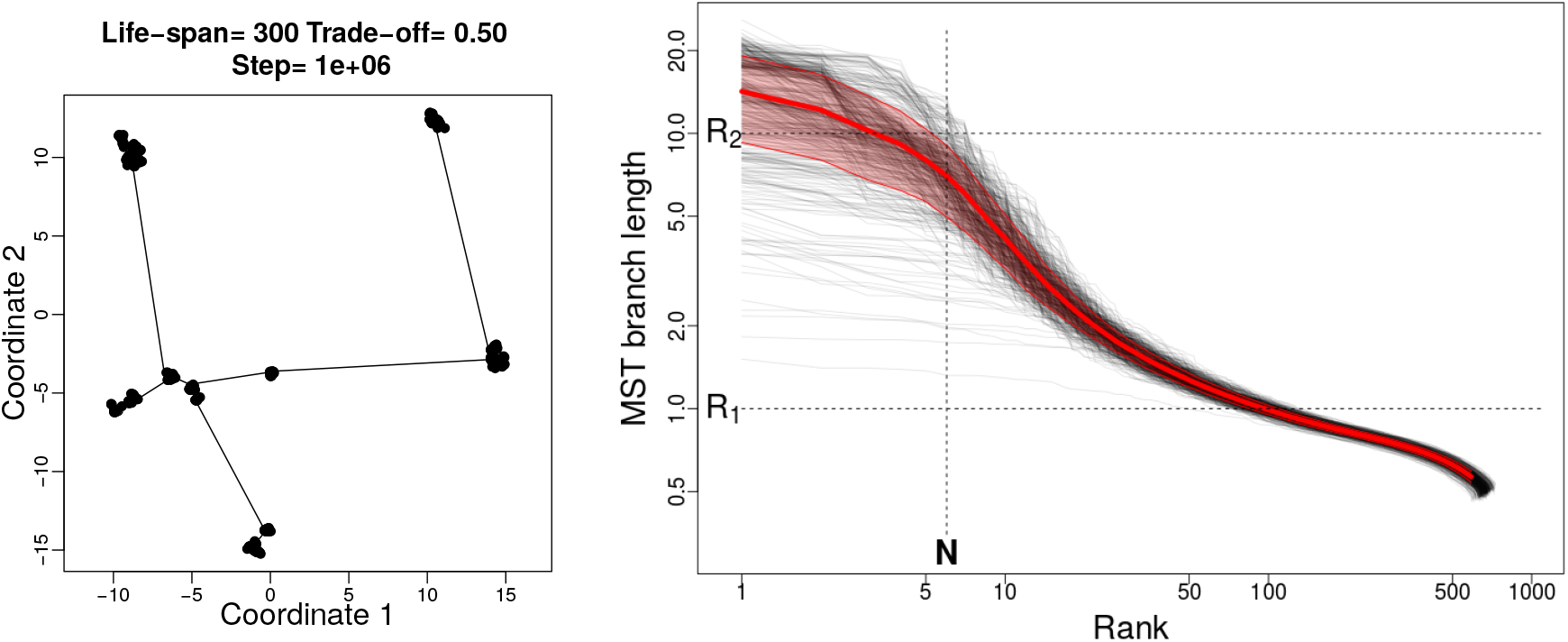
Left: 2D representation of minimum spanning tree (MST) of a typical snapshot of the simulation for *δ = 0.5, λ = 300, μ = 0.001 and m = 0.02* The community of this snapshot consists of 667 strains. Right: gray curves show the sorted length of the edges of MST versus their ranks for 500 snapshots of a system with the aforementioned parameters. Red curve shows the average of all curves and the red shaded area show the standard deviation of them.

**Figure 5.**
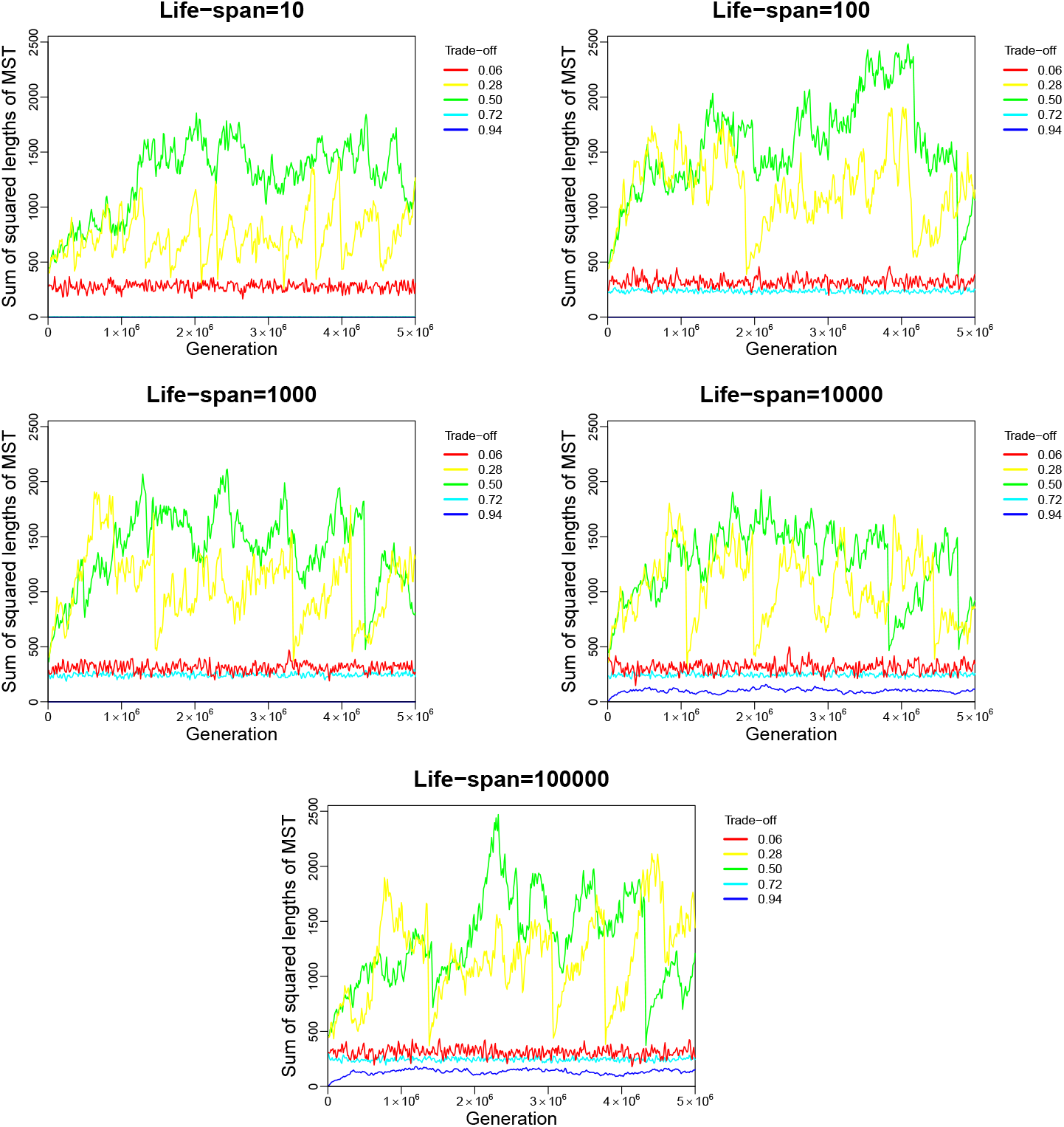
Each plot shows how SMST changes over generations for simulations with fixed lifespans but different trade-offs. For very strong trade-offs diverse strategies can not be adopted. For very weak tradeoffs diverse strategies can emerge easily but among them extreme strategies (Darwinian Daemons) very fast dominate and diversity decreases. Sustainable diversity emerges in moderate trade-offs. Very short lifespans prevent increase in diversity, specially for strong trade-offs. Simulations are done with *μ = 0.001 and m = 0.02*.

**Figure 6.**
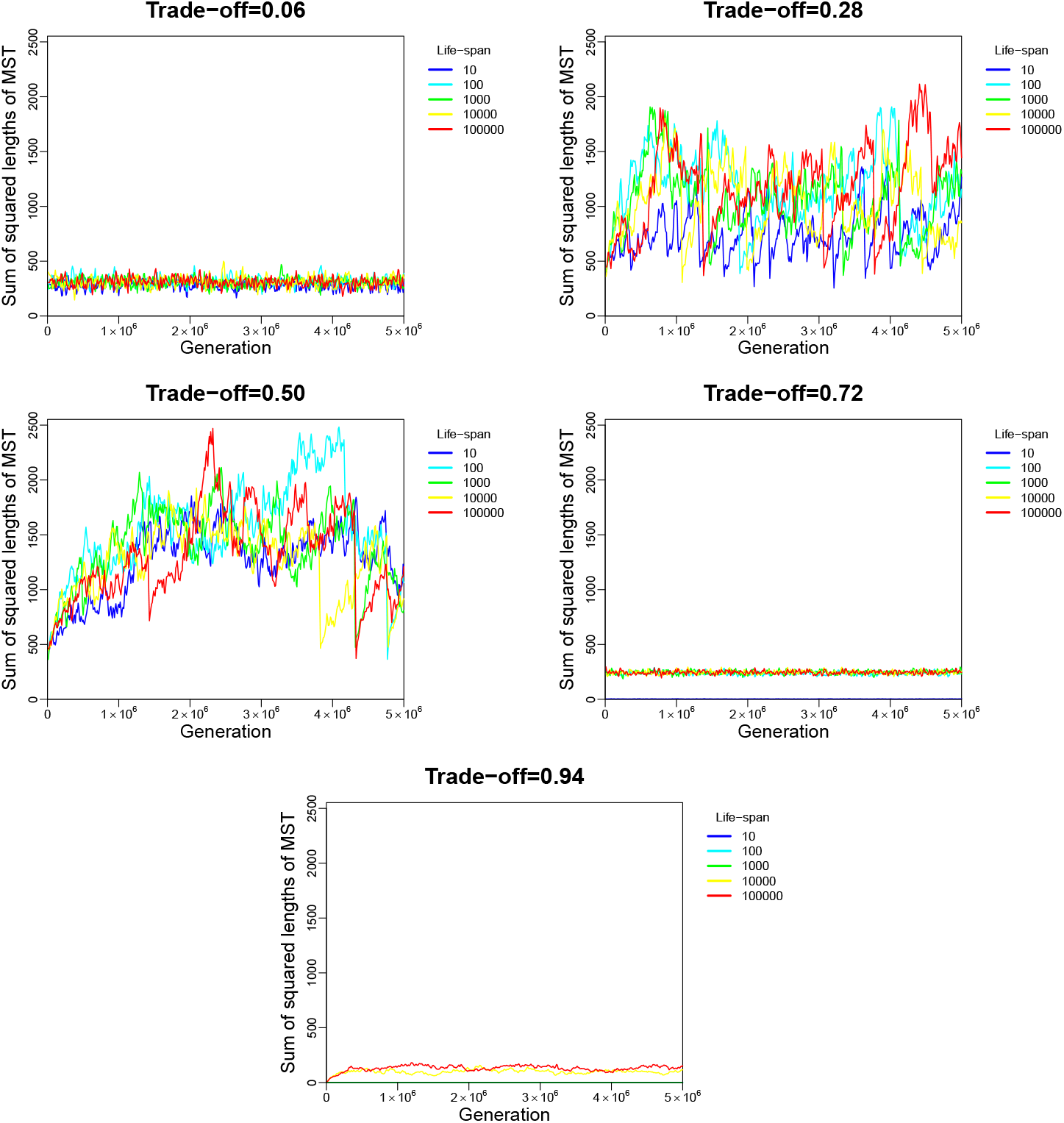
Each plot shows how SMST changes over generations for simulations with fixed trade-off but different lifespans. lifespan has not a big effect for weak and moderate trade-offs but for strong trade-offs, short lifespans yield to complete mass extinction. Simulations are done with *μ = 0.001 and m = 0.02*.

**Figure 7.**
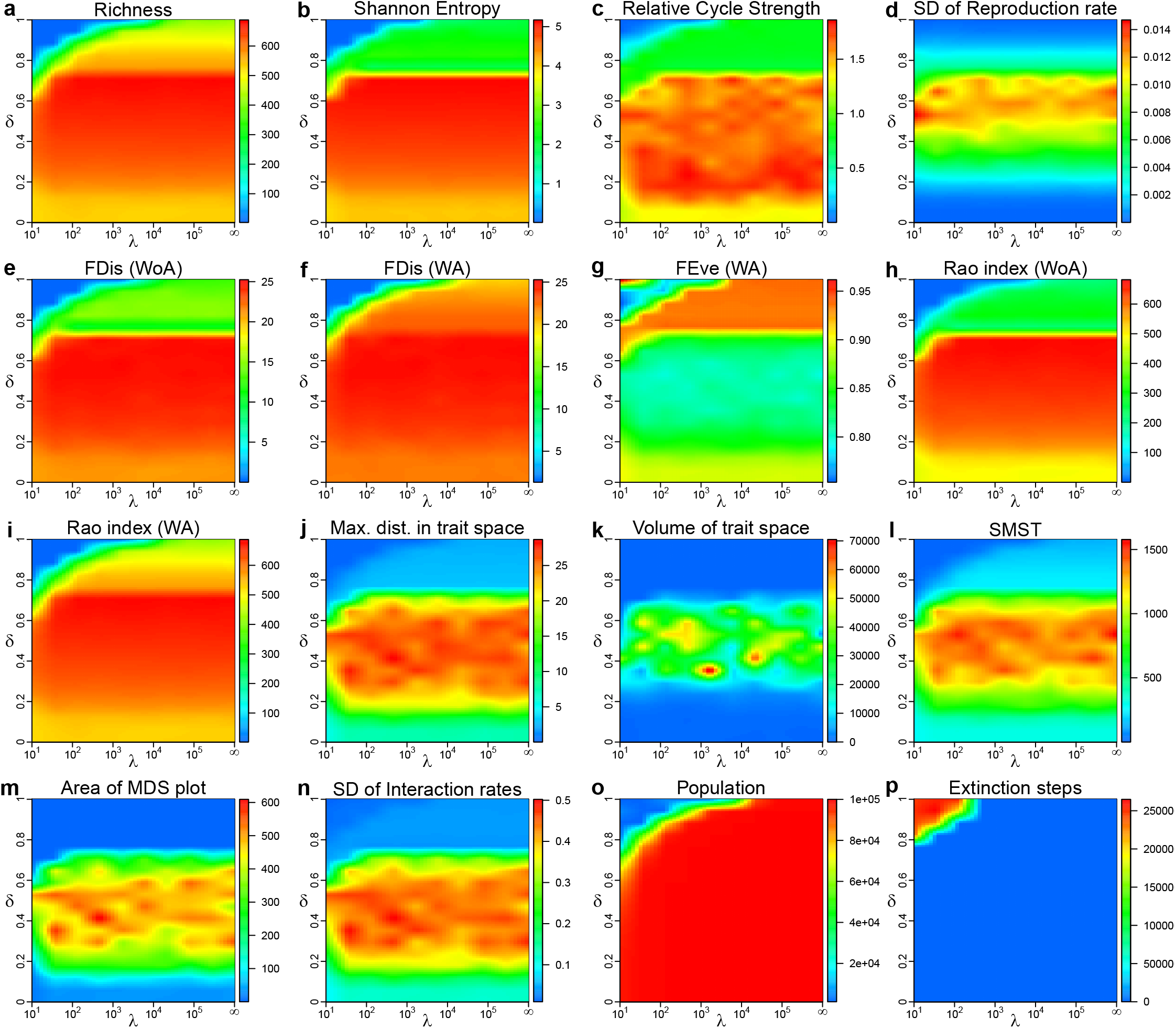
A) **Richness**: Number of different strains in community. b) **Shannon entropy**: which is a measure of evenness in strain population. c) **Relative Strength of cycles** of size 3 compared to random networks. (see SI-6) d) **Standard deviation of reproduction rates** as a measure of diversity in reproduction strategy. e) **Functional dispersion** (FDis, without using the abundance vector): measures the mean distance of individual strains in trait space to their centroid (dbFD function in FD package, version 1.0-12, in R) f) **Functional dispersion** (FDis, using the abundance vector): measures the mean distance of individual strains (weighted by abundance vector) in trait space to their centroid (dbFD function in FD package, version 1.0-12, in R) g) **Functional evenness** (FEve, using the abundance vector): quantifies functional evenness and is higher when species are spread homogeneously in trait space. When disruptive selection produces colonies of localized strains in trait space this index decreases (dbFD function in FD package, version 1.0-12, in R). h) **Rao’s quadratic entropy** (without using the abundance vector): measures mean functional distance between two randomly chosen individuals (dbFD function in FD package, version 1.0-12, in R). i) **Rao’s quadratic entropy** (using the abundance vector): measures mean functional distance between two randomly chosen individuals (dbFD function in FD package, version 1.0-12, in R). j) **Maximum distance in trait space** between strains. k) **Volume of trait space**: calculated by multiplication of eigenvalues of factor analysis. l) **Sum of squared length of minimum spanning tree in trait space** (SMST). m) **Area of MDS plot**: calculated by multiplication of the two first eigenvalues of factor analysis. n) **Standard deviation of interaction rates**. o) **Community population**: Number of Individuals in community. p) **Number of mass extinction events** over 5 × 10^6^ generations. Each plot is the average of corresponding index over 3 simulations each over 5 × 10^6^generations. Simulations are done with *μ = 0.001, m = 0.02* and *N_s_ = 10^5^*.

### VIII Size of the system

Simulations with *N_s_* = 1 × 10^4^, 3 × 10^4^,1 × 10^5^, 3 × 10^5^, λ =∞ to and a set of trade-off parameters (0 ≤ δ *<* 1) were done to check the effect of size of the system. One of the diversity indexes (SMST) is plotted in Fig. 8 for different sizes. As it is clear by increasing *N_s_* the final diversity in trait space is higher and the drop in δ ≈ 0.7 is sharper which is typical of a phase transition.

**Figure 8.**
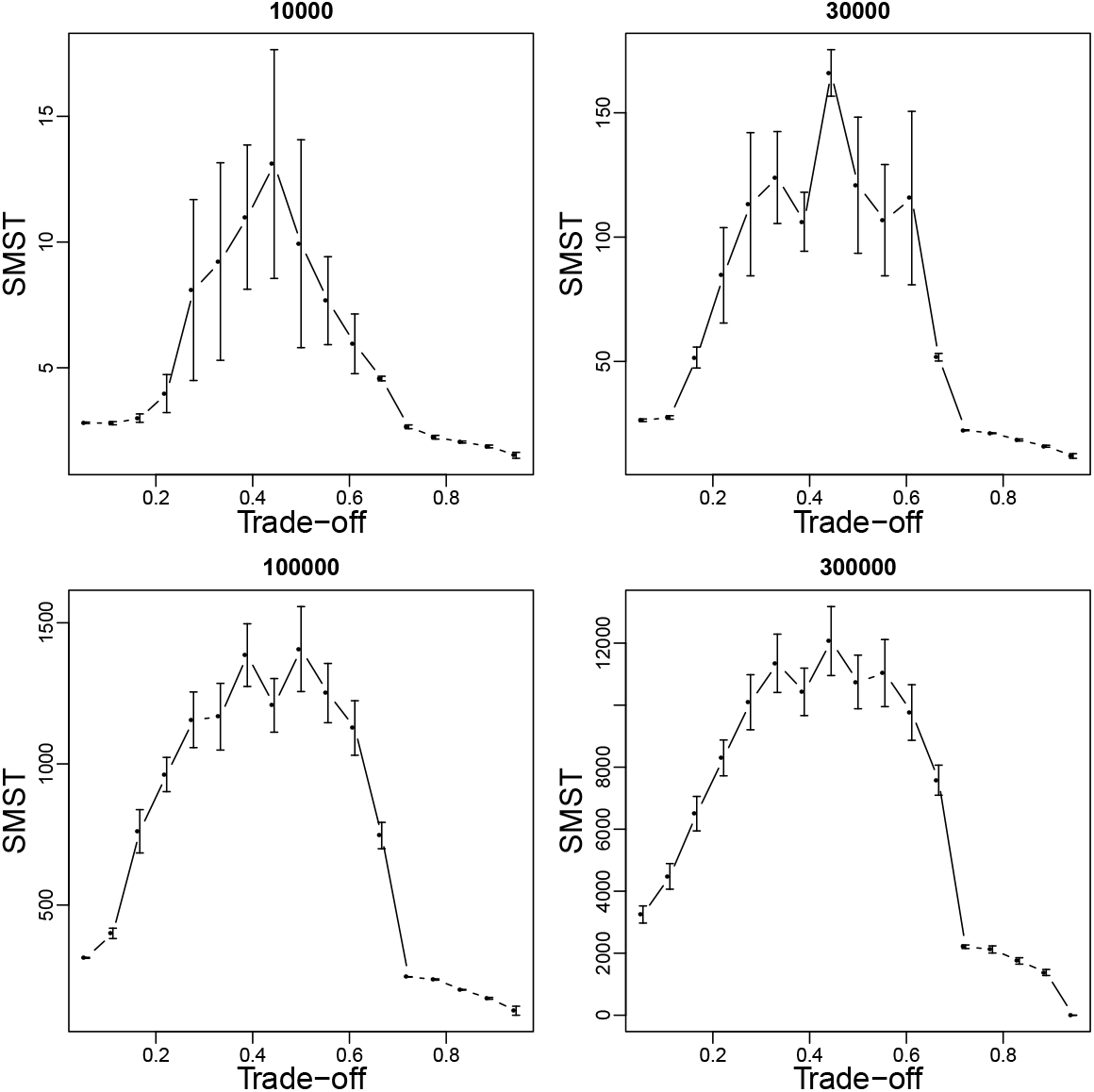
Functional diversity as measured by size of minimum spanning tree (SMST) as function of trade-off parameter for different system sizes *N_s_*. Simulations are done with *λ= ∞, μ = 0:001 and m = 0:02*.

### IX Frequency-dependent selection

Frequency-dependent selection mediated by interaction of species could be a source for temporal correlation between eco-evolutionary events, e.g. speciation, invasion and extinction of species. To examine if there is such a correlation we used the distribution of inter-event times, i.e. distribution of intervals between occurrence of events. For a completely random process (Poisson process) this distribution follows an exponential distribution and deviation from exponential is a signature of correlation between events. Fig 9 shows interevent distribution of ITEEM data and compares it with the best fit of geometric distribution (discrete version of exponential distribution) to data. The clear deviation from Poisson process shows that speciation and extinctions are not just random events but after occurrence of an event, with a delay (≈ 10000 generations), the probability of observing a new event is higher than a random process.

**Figure 9.**
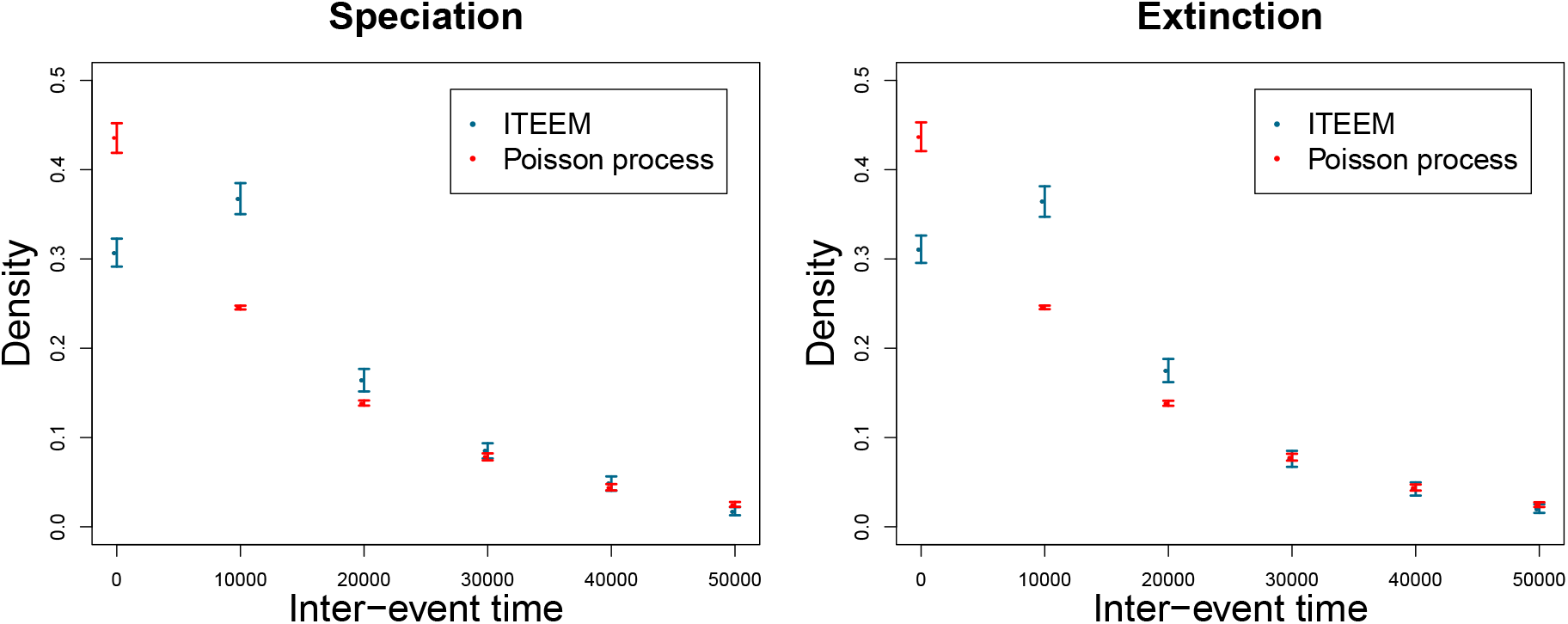
Distribution of time interval between speciation events (left) and extinction events (right). Blue points and error bars: data from 24 ITEEM simulations, each of 5 × 10^6^ generations (*δ = 0.5, λ = 10^5^ … ∞ N_S_ = 10^5^, μ = 0.001, m = 0.02*). Error bars are ±2 standard deviations calculated by bootstrapping. Red points and error bars: maximum likelihood fit (function fitdistr in R-package MASS, version 7.3-44) of the simulated data to a geometric distribution (discrete version of an exponential distribution), corresponding to an assumed Poisson process. Error bars are ±2 standard deviations estimated by the maximum likelihood fit.

### X Mutation Rate

Fig. 10 show behavior of typical diversity index (SMST) over δ and λ for different mutation rates. Although the overall dependency on trade-off strength and lifespan is the same for a wide range of mutation rates but the value of diversity indexes depend on mutation rate: the smaller the mutation rate the lower the diversity in community.

**Figure 10.**
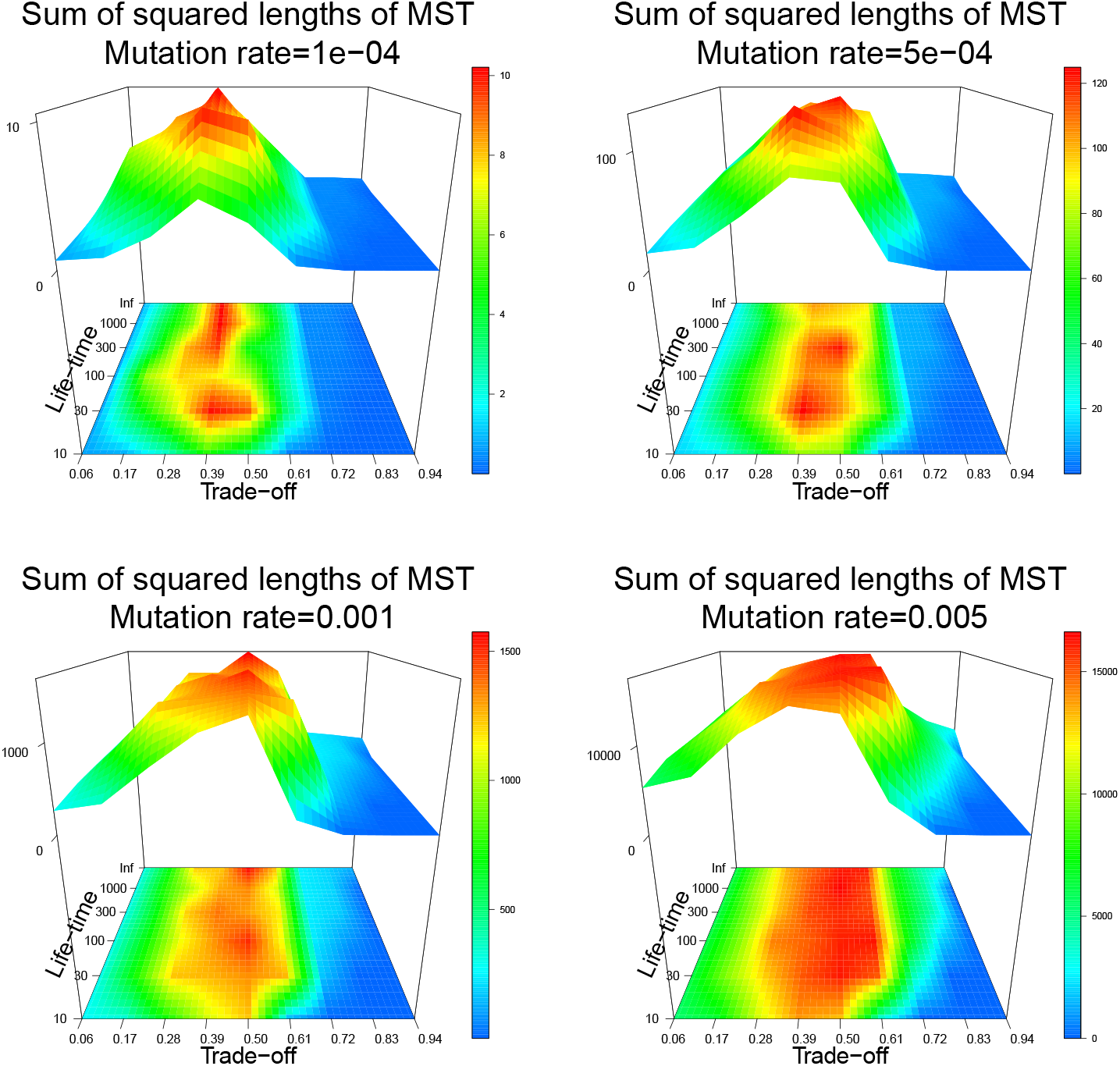
SMST for different mutation rates versus trade-off and lifespan.

**Figure 11.**
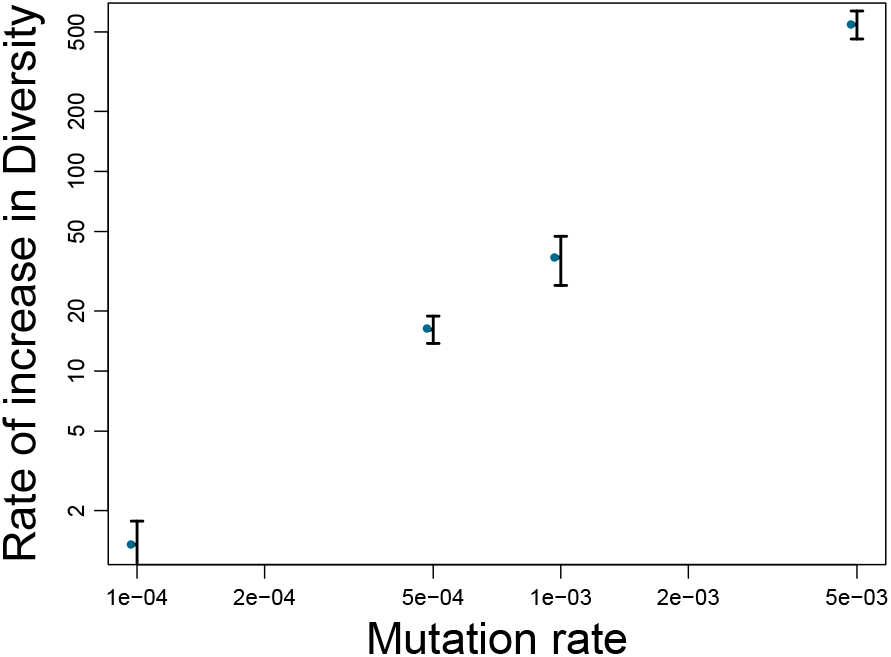
Rate of increase in diversity (measured as increase of SMST per 10000 generations) for different mutation rates. Rate of increase in diversity is calculated by fitting a line to the first 80000 generations of each simulation and averaging is over 5 different simulations. Error bars show the errors estimated by fit. Note the logarithmic scale of the plot.

### XI Neutral Model

To compare the diversity generated by genetic drift (neutral model) with diversity generated under selection pressure induced by competition with moderate trade-offs, we do simulations in which all species compete equally for resources, i.e. ***I**_αβ_ = 0.5* for all pairs of species, but traits evolve by mutation as before. In the neutral model reproduction rates should also be the same for all species so we attribute the same reproduction rate in each simulation to all species. We carry out simulations for reproduction rates *r* = 0.1, 0.5, 0.9.

Distribution of strains over trait space (Fig. 12) show that genetic drift is able to spread trait vectors in trait space and produce cloud of strains but the size of diversity generated by it is much smaller than that of communities evolved under biotic selection pressure mediated by competition under moderate trade-offs.

**Figure 12.**
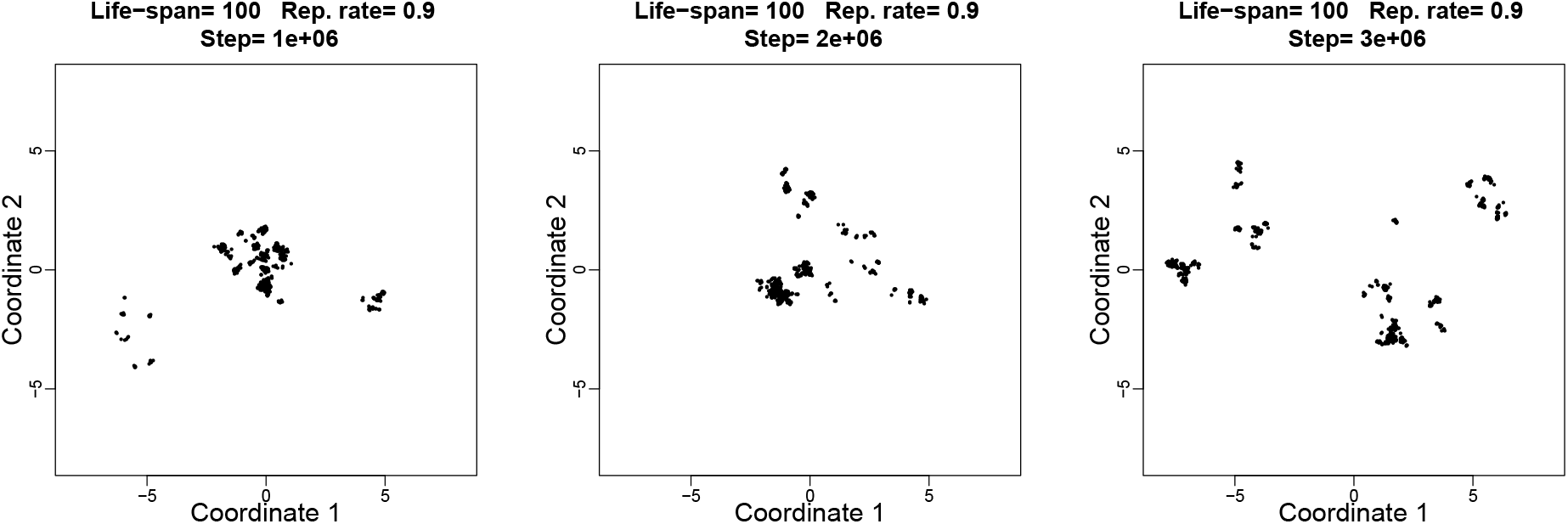
Trait space of a community evolved under neutral model in 3 time step: 1 × 10^6^,2 × 10^6^ and 3 × 10^6^ with *λ = 100, r = 0.9, μ = 0.001 and m = 0.02*. Note the small size of the trait space in comparison to a non-neutral model (SI Fig 3).

In order to show clearly the difference between the diversity produced in both models we also studied the diversity measures and other parameters. In Fig. 13, left panel in each row shows changes of a typical diversity index (SMST) over time for a simulation of genetic drift for one lifespan (with 3 different reproduction rates) and compares the results with a typical simulation with competitive selection pressure for a moderate tradeoff (δ= 0.56). Right panels illustrate the corresponding plots but for cycle formation. Relative strength of cycles in these simulations has big fluctuations around a value less than one, without any stable pattern over time. This shows that, as it is expected, dynamics of community in the neutral model is determined by fluctuations.

**Figure 13.**
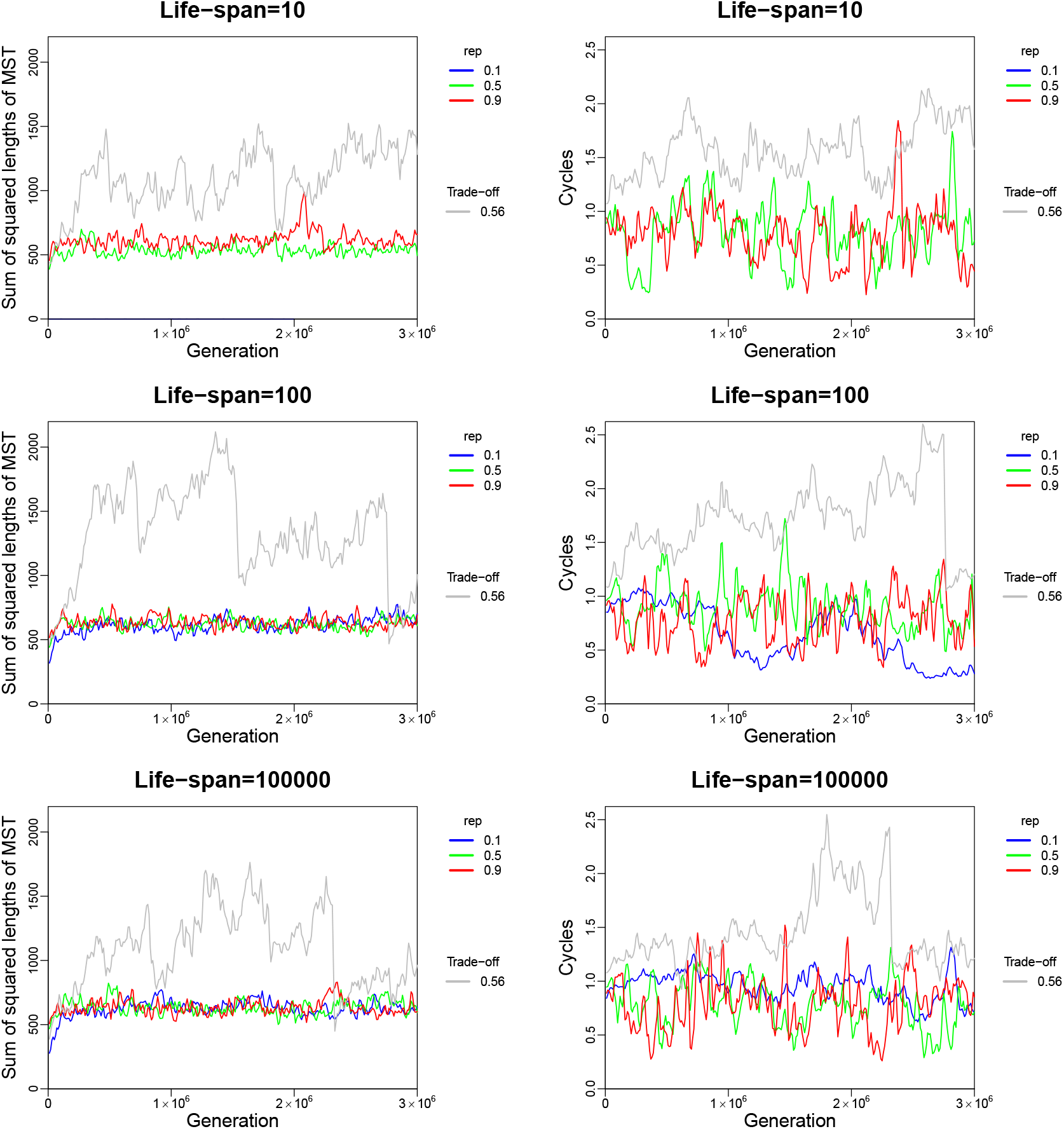
Plots in the left and right panels show how SMST and relative strength of cycles, respectively, change over generations for 3 different lifespans. Colored curves are the results of the neutral model with different reproduction rates (*r = 0.1, 0.5,0, 9*). The results are compared with the outcome of one simulation with the corresponding lifespans and trade-off strength of 0.56 (gray curves). For *λ =10 and r = 0.1* population goes to extinction very fast.

